# Adversarial training improves model interpretability in single-cell RNA-seq analysis

**DOI:** 10.1101/2023.05.17.541170

**Authors:** Mehrshad Sadria, Anita Layton, Gary D. Bader

## Abstract

For predictive computational models to be considered reliable in crucial areas such as biology and medicine, it is essential for them to be accurate, robust, and interpretable. A sufficiently robust model should not have its output affected significantly by a slight change in the input. Also, these models should be able to explain how a decision is made. Efforts have been made to improve the robustness and interpretability of these models as independent challenges, however, the effect of robustness and interpretability on each other is poorly understood. Here, we show that predicting cell type based on single-cell RNA-seq data is more robust by adversarially training a deep learning model. Surprisingly, we find this also leads to improved model interpretability, as measured by identifying genes important for classification. We believe that adversarial training will be generally useful to improve deep learning robustness and interpretability, thereby facilitating biological discovery.

## Introduction

Deep learning is important in processing big biological data structures and has reshaped our ability to analyze large-scale datasets, such as single-cell data (1). However, the traditional black-box nature of deep neural networks (DNNs) remains a major obstacle to their wide adoption in applications where mechanistic insight is important (2). A number of interpretability techniques have been developed to comprehend DNNs (3–5). For example, by using these techniques we can identify input features that significantly impact the final classification. Gradient calculations are also frequently employed to provide each input feature with a weighted significance score that reflects the impact of that feature on model predictions (6). Deep learning models can also have problems with robustness, where their predictions are highly sensitive to small perturbations in input data (7). These small perturbations can be specifically generated based on the trained model, in which case they are called adversarial data (8,9). Adversarial training is a recent advance in deep learning, so far mainly applied with image and text input, that results in more robust and generalizable models (10,11). Adversarial training involves supplementing training data with generated adversarial instances during each training loop which leads to more robust models. For image applications, adversarially trained DNNs have been observed to yield loss gradients that are visually more interpretable than those from analogous models without adversarial training (12,13), but the interplay between robustness and interpretability in deep learning is poorly understood.

To study the relationship between robustness and interpretability in computational classification, we analyzed the task of cell-type classification based on single-cell RNA-seq data. The human body is made up of trillions of cells of many different cell types. Single-cell genomics technology has made it possible to measure gene expression profiles at single-cell resolution, providing an unprecedented opportunity to study the processes of multicellular organism growth, as well as disease and treatment response (14). To study these processes, cell types and states must be reliably identified to observe how they change over time (15). We also need to know key genes, such as gene expression markers or master regulators, that are useful for cell type identification and mechanistic understanding of the underlying biology (16).

In this work, we explored the effect of using adversarial training on the performance (accuracy, robustness, and interpretability) of deep learning models trained for cell-type classification using a range of simulated and real single-cell RNA sequencing data. We used several interpretability techniques to identify genes that are essential for cell type classification. Interestingly, we find that adversarial training increases both the robustness and interpretability of the resulting models and can be used to discover new biological insights.

## Results

### Adversarial training improves robustness for cell type classification

To explore model robustness using adversarial training, we selected single-cell classification as an example task. A multi-layer perceptron architecture (the detailed architecture in Supplementary Table 1) was selected to implement this task. Single-cell RNA-seq data, represented as a cell by gene matrix, with a given set of ground truth cell classes is used as input. Initially, we use simulated data generated by SERGIO (17) to ensure we work with a concrete ground truth data set. SERGIO enables users to specify the number of cell types to be simulated, given a simulated gene regulatory network (Supplementary Figure 1). We used SERGIO to generate a gene expression matrix of 2700 cells and 1200 genes with 9 cell types. The gene-by-cell matrix and cell type labels were used to train a classifier with a hyperparameter search on the number of layers and nodes. The classifier achieves an accuracy of 98.2% on the simulated data. We then used two established methods to generate adversarial data which is required for adversarial training: Projected Gradient Descent (PGD) and Fast Gradient Signed Method (FGSM) (8,18). These take the trained model and introduce noise in the input data in the direction of the model gradient that has the greatest impact on the model’s accuracy. However, we must tune the amount of noise, ε, in this procedure, as adding too much or not enough noise will not result in useful adversarial data for training. In typical applications such as computer vision, the value of ε is well-established for different methods (8,18). However, tabular data, like scRNA-seq, has received less attention in identifying the appropriate ε value. A good ε value is one that causes a noticeable reduction in the classifier’s accuracy while not substantially interfering with the structure of the input data. Therefore, we varied the ε value while generating adversarial training data using PGD and FGSM with our simulated gene expression data. The newly generated adversarial data is combined with the original data to create an adversarial training data set. We evaluate the ε value in two ways: classification performance of the original model on the adversarial training data set and evaluation of the stability of the global data structure using UMAP visualization (Fig.1). Values of ε between 1 - 1.2 significantly decreased classifier accuracy while maintaining global structure when the model is subjected to FGSM perturbation (Fig. 1b and 1d blue lines). However, larger ε values (e.g. 3.2) cause the global structure to degrade, indicating that the selected value is too large (Fig. 1c). We next test the effect of adversarial training using our established ε value, comparing model accuracy without adversarial training (Fig. 1d,e blue lines) to accuracy with adversarial training (Fig. 1d,e orange lines). We find that training with adversarial data using both FGSM and PGD significantly strengthens the model’s robustness, raising accuracy from as low as 30% with standard training to almost perfect 100% accuracy with adversarial training.

**Figure 1.**
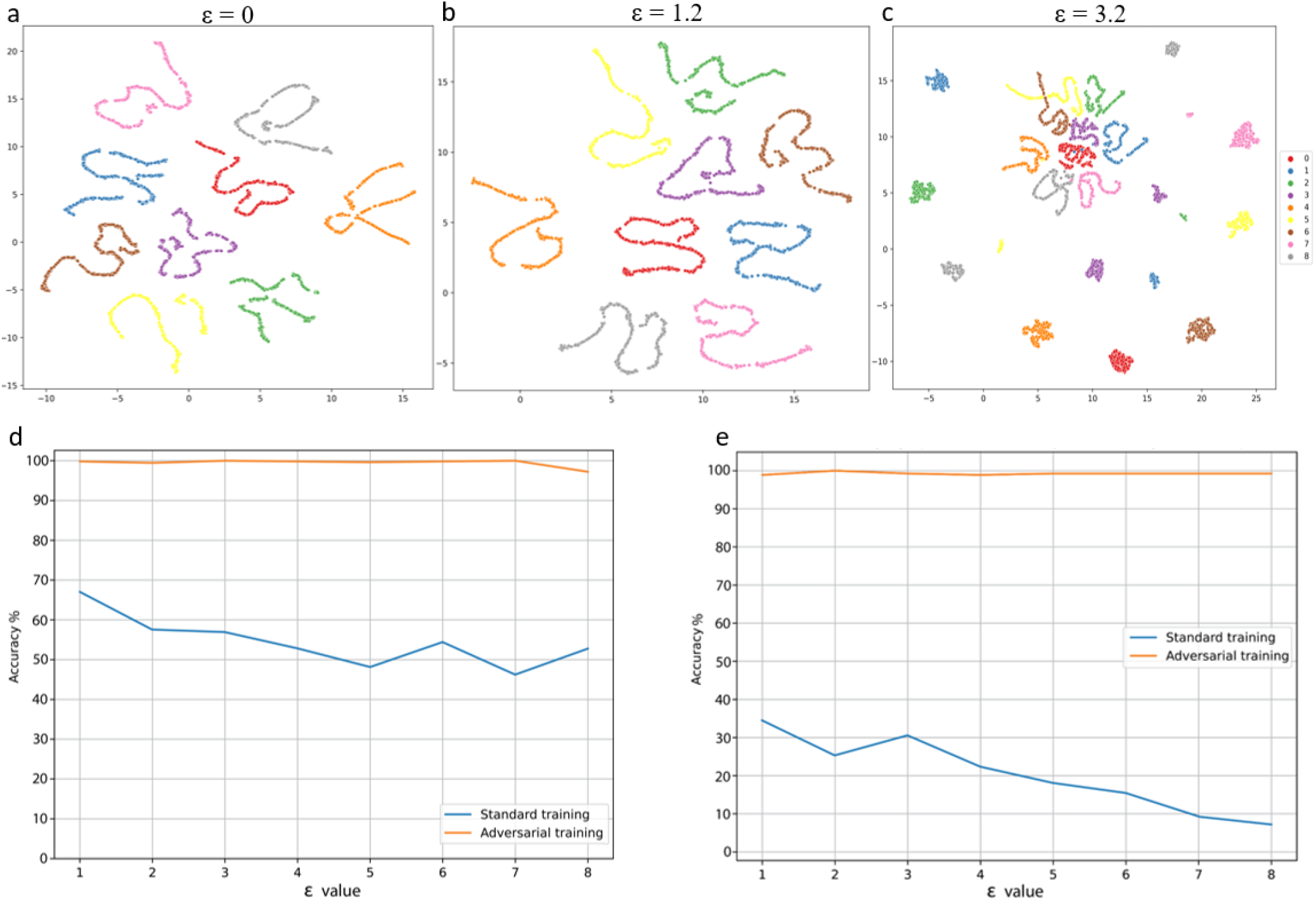
Effects of adversarial attack on data structure and model accuracy. UMAP plots (a) of original data, (b) with an adversarial attack using FGSM with an epsilon value of 1.2, which preserves data structure, and (c) with an adversarial attack with a larger epsilon value of 3.2, which substantially changes data structure. Effect on model accuracy following FGSM (d) and PGD attacks (e). Adversarial attacks reduce model accuracy substantially (blue lines), but with appropriate training, high accuracy can be achieved (orange lines).

### Adversarial training improves model interpretability

While adversarial training fortifies a neural network against adversarial perturbations and increases the robustness of the model, its effects on interpretability are not well studied. Many DNN interpretability methods are based on analyzing weights and gradient loss, therefore, adversarial training may affect model interpretability (6). To study this relationship, we applied an adversarially trained neural network to our SERGIO-simulated scRNA-seq data generated to include 65 predefined key genes to classify cell-types and identified significant genes for each cell type using six different DNN interpretability methods. These six methods are saliency maps (3), activation maximization (19), Local Interpretable Model-agnostic Explanations (LIME) (20), and three variants of SHapley Additive exPlanations (SHAP) (21): gradient explainer, deep explainer, and kernel explainer. All these methods use a local linear approximation to identify important features of a model. However, the loss functions and local neighbourhood definition differ among these methods, which often results in discrepancies and differences in their results.

To measure the effect of adversarial training on the accuracy of the interpretability methods, we compute the number of predefined key genes correctly detected by our interpretability methods for each cell type with and without adversarial training. For most of the cell types, an adversarially trained neural network was able to detect more key genes than the non-adversarially trained model (Fig. 2a-d and Supplementary Figure 2). These results indicate that, in general, adversarial training improves the accuracy of the interpretability methods. As noted by the no-free-lunch theorem for explanation methods, there is substantial variability in the performance of various interpretability methods (Supplementary Figure 3), which depends on the characteristics of the data and the local context (22). Based on this observation, we use a voting mechanism that considers the aggregate of significant genes recognized by several interpretability methods and calculates a “consensus importance score”. This score denotes the number of interpretability methods that identify a specific gene as one of the top N genes for a given cell type. Specifically, we consider the top 20 most important genes identified by each method and rank the genes based on the majority vote. Results in Fig. 2e show that adversarial training increases the number of correctly identified key genes in almost all cell types for this majority vote approach. These results suggest that adversarial training not only can improve model robustness but also its interpretability, as measured by the number of correctly identified key genes.

**Figure 2.**
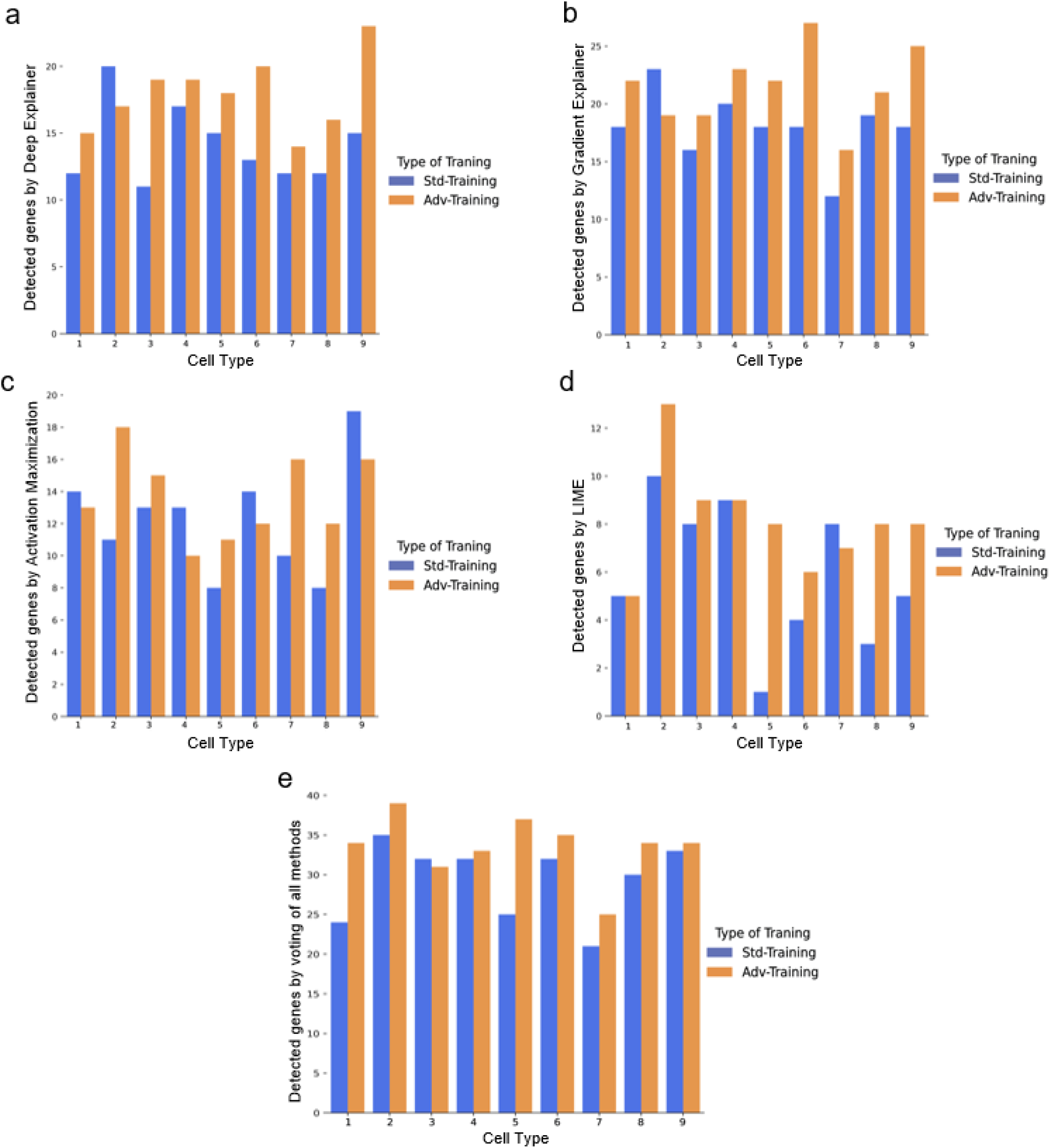
The effect of adversarial training on the model’s interpretability. a, the results of using the Deep Explainer b, Gradient Explainer, c Activation maximization, d, LIME, and e, the number of detected key genes using an aggregate result of all methods before and after adversarial training.

### Using adversarial training can discover key genes and pathways in single-cell RNA-seq data

To establish the applicability of the adversarial training in real biological data, we analyze a mouse hippocampus development scRNA-seq dataset of 18231 cells, 14 cell types, and 3,001 genes (23) (Fig. 3a). We first train our non-adversarially-trained DNN classifier to predict known cell types. The classifier performs well for the majority of cell types, except for the distinction between mature and immature granulocytes (Fig. 3b). When the model is adversarially attacked using PGD, its accuracy decreases significantly (Supplementary Figure 4). To increase its robustness, we adversarially train the classifier, apply all six interpretability methods to the resulting model to identify key genes for each cell type (Supplementary Figure 5), and compute the “consensus importance score” for each gene-cell type pair (N = top 150 genes). We confirm that adversarial training of the model using both FGSM and PGD significantly improves its robustness (Supplementary Figure 4a,b). We then clustered the genes according to their consensus importance scores and visualized the results as a heatmap (Fig. 3c). Notably, genes located on the left side of the heatmap are selected as important for classifying the majority of cell types. In contrast, genes located on the right side of the heatmap are important for classifying specific types. Table 1 shows the top predicted important genes based on the consensus importance scores computed for each cell type. Most genes in this table (50 of 59) are known to play a key role in that specific cell type or in neuronal development in general (24–28) (Supplementary Table 2 contains genes and refrences to the previously reported studies).

**Figure 3.**
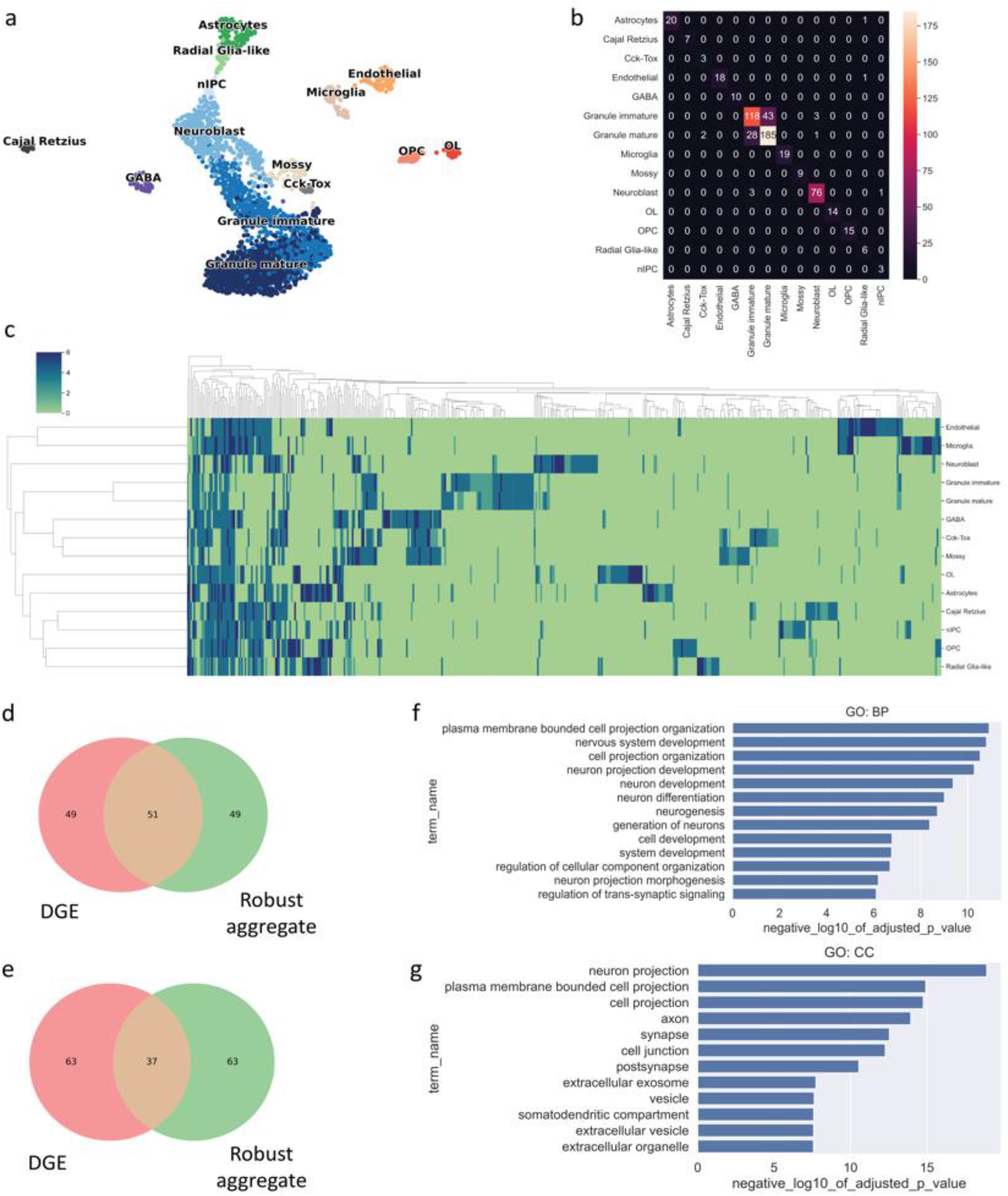
Applying adversarial training to identify important genes in mouse hippocampus development. a, UMAP visualization of mouse hippocampus development data (14 cell types). b, the confusion matrix of the classifier. c, the consensus importance scores of genes determined by applying multiple interpretability methods visualized as a clustered heatmap. (d,e), Venn diagrams show the comparison of the number of genes detected by differential gene expression and adversarial training based on importance prediction for astrocyte and endothelial cells, respectively. (f, g) Show the pathway and cellular component gene set enrichment analysis, respectively, of the predicted important genes. DGE = differential gene expression.

**Table 1.** Genes predicted as important (high consensus importance score) for each cell-type for the mouse hippocampus dataset. The bolded genes are the ones that were not detected by DGE but were detected by consensus importance score.

| Cell type | Predicted important genes |
| --- | --- |
| Astrocytes | Gad2, Dbi, Fabp7, Igfbpl1, Cplx2 |
| Cajal Retzius | Dbi, Itm2b, Tmsb10, Cplx2, **Atp5g1** |
| Cck_Tox | Gad1, Igfbpl1, **Scg5, Ppfia2, Lamtor2** |
| Endothelial | Sept4, Klf2, Igfbpl1, Snca, Cnih2 |
| GABA | Nrgn, Npy, Itm2a, Cst3, Calm2 |
| Granule immature | Ncdn, **Lin7b**, Camk4, Snca, **Tspan3** |
| Granule mature | **Dmtn, Rbm25, Rap1b**, Cpe, C1ql3 |
| Microglia | Nrgn, Hexb, P2ry12, Cnih2, Cplx2 |
| Mossy | B2m, Trappc4, Camk4, **Cntn1, Mycbp2** |
| Neuroblast | **Dnaja1, Arpc1a**, Cfdp1, **Nell2, Prdx2** |
| OL | Camk2a, Mllt11, Sept7, Tubb4a, **Fez1** |
| OPC | Stmn1, Stmn4, Fabp7, Scrg1, Golga7 |
| Radial Glia-like | Dbi, Zbtb20, Slc1a2, Scrg1, Ncdn |
| nIPC | Camk2b, Bcl11b, **Aplp1**, Arl6ip1, **Hmgn3** |

To further validate the important genes we predict, we perform pathway and cellular location enrichment analysis on the 15 most important predicted genes from each cell type (total = 210). This analysis shows that pathways related to nervous system development and neurogenesis are significantly enriched in these genes (Fig. 3f), as well as brain-related cellular compartments such as neuron projection, synapse, and axon (Fig. 3g). As a control, we compare our predicted important genes to the list of differentially expressed genes, selected as a standard method to identify important genes. We compute for astrocytes and endothelial cells the top 100 predicted important genes by our model and by differential gene expression analysis. For astrocytes, half of the top genes are shared between our method and differential gene expression analysis; for endothelial cells, about one-third (see Fig. 3d,e). Examining the top five genes from each cell type, we observe that (50 of 59 genes) are only found by our consensus importance score, and most of these are known to be important for the brain in the literature (Supplementary Table 2). Thus, our method identifies important genes not identified by the standard method.

As an additional test, we repeated our analysis on a mouse pancreas scRNA-seq dataset collected from the 15.5th day of embryonic development (29) with 2531 cells clustered in seven cell types (Fig.4). By using the adversarially trained model using PGD and FGSM, we were able to increase the model robustness (Fig. 4d,e). We again see that many of the important genes predicted by our method play a key role in pancreas development (Table 2) (30–36) and strong pathway and cellular component enrichment in the genes for each cell type related to pancreas functions (regulating cell secretion, hormone activity, hormone transport, and pancreas development).

**Figure 4.**
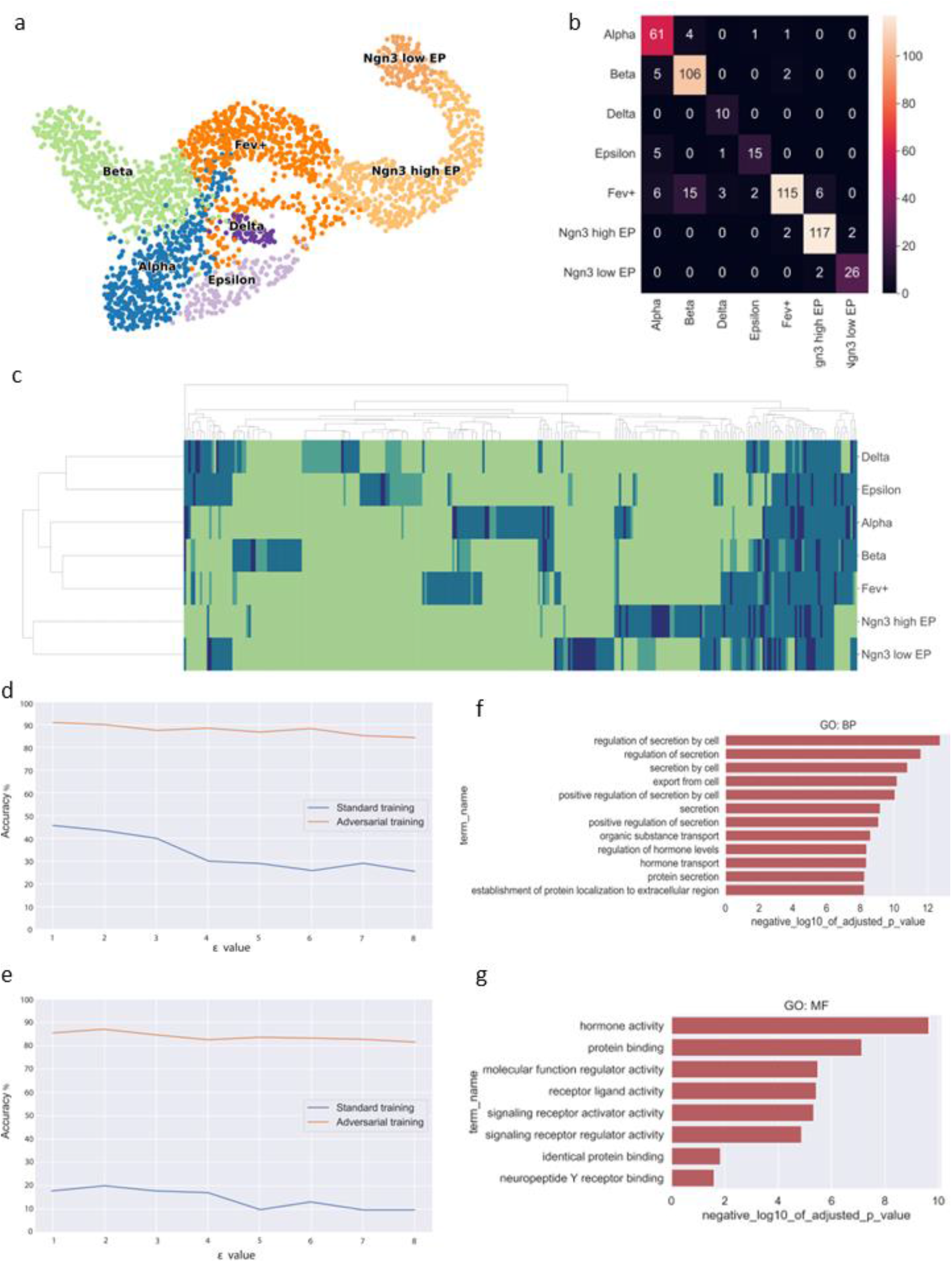
A Representation of the mouse pancreas data after applying interpretability methods to an adversarially trained classifier. a, The UMAP visualization of the data, highlighting the clustering of different cell types. b, The confusion matrix of the classifier, shows the model’s performance in cell-type classification. c, The heatmap of the consensus importance scores was determined by applying multiple interpretability methods for each gene cell-type pair. d,e, The accuracy of the model before and after adversarial training using PGD and FGSM methods respectively. Lastly, f and g, showcase the gene set enrichment analysis of the crucial genes identified by the consensus importance scores, providing a deeper understanding of the underlying biological process and molecular function.

**Table 2.**
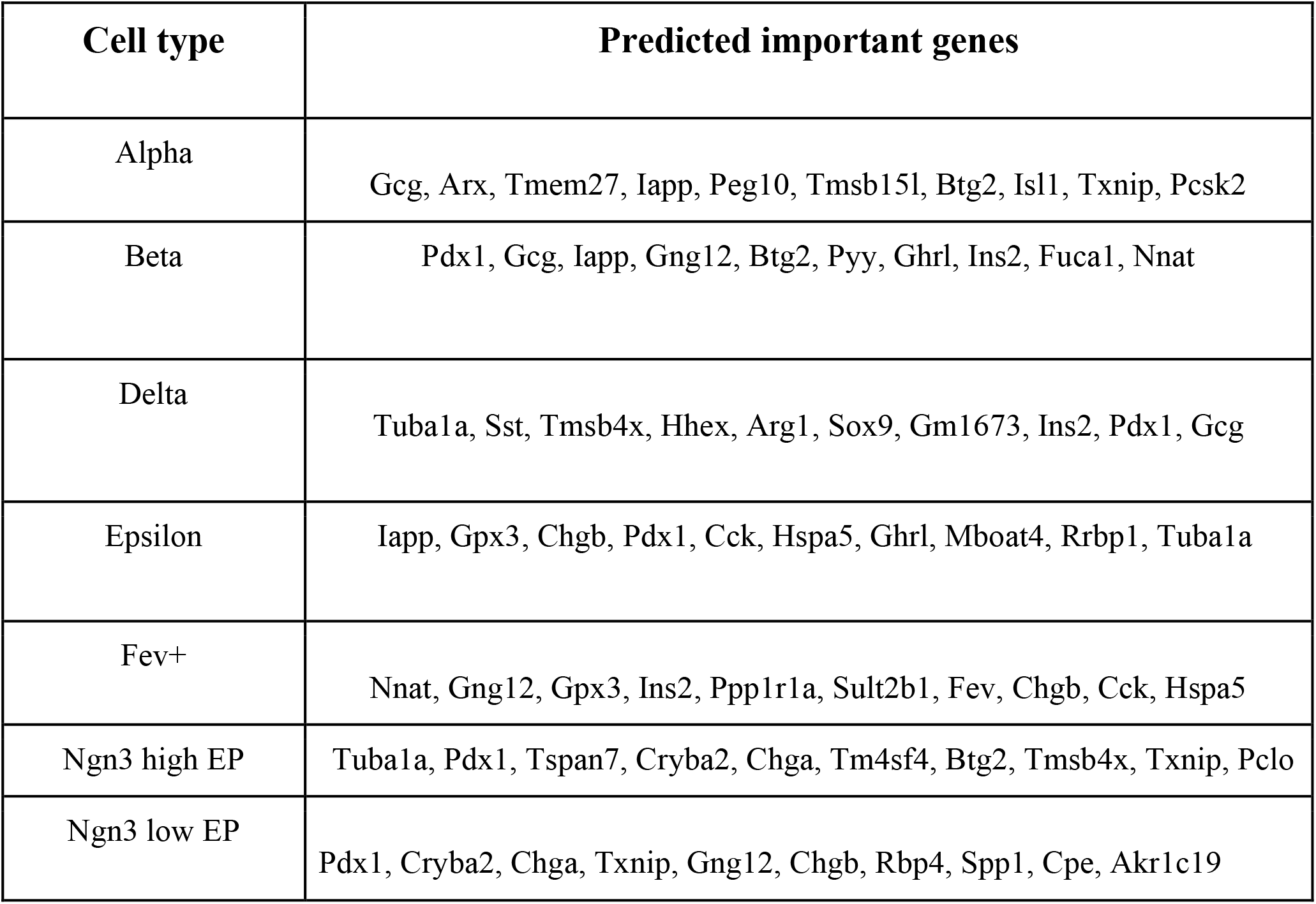
Gene candidates detected for each cell-type using consensus scores for the developing mouse pancreas data. The table includes columns for the cell-type and the detected gene candidates.

| Cell type | Predicted important genes |
| --- | --- |
| Alpha | Gcg, Arx, Tmem27, Iapp, Peg10, Tmsb15l, Btg2, Isl1, Txnip, Pcsk2 |
| Beta | Pdx1, Gcg, Iapp, Gng12, Btg2, Pyy, Ghrl, Ins2, Fuca1, Nnat |
| Delta | Tuba1a, Sst, Tmsb4x, Hhex, Arg1, Sox9, Gm1673, Ins2, Pdx1, Gcg |
| Epsilon | Iapp, Gpx3, Chgb, Pdx1, Cck, Hspa5, Ghrl, Mboat4, Rrbp1, Tuba1a |
| Fev+ | Nnat, Gng12, Gpx3, Ins2, Ppp1r1a, Sult2b1, Fev, Chgb, Cck, Hspa5 |
| Ngn3 high EP | Tuba1a, Pdx1, Tspan7, Cryba2, Chga, Tm4sf4, Btg2, Tmsb4x, Txnip, Pclo |
| Ngn3 low EP | Pdx1, Cryba2, Chga, Txnip, Gng12, Chgb, Rbp4, Spp1, Cpe, Akr1c19 |

## Discussion

The recent rapid developments in single-cell technology have enabled the analysis of biological data at unprecedented scale and resolution. In addition to transcriptomics data, single-cell technology includes more than 200 techniques that profile the genomic, epigenetic, and proteomic data in individual cells (37). By providing detailed molecular information on cells, single-cell atlases can facilitate the identification of key genes and pathways linked to healthy and disease states, patient classification, and the development of targeted therapies (29,38–40). It’s essential for machine learning models used in analyzing biological and clinical data to be accurate and robust. Interpretability is also crucial as it provides insight into the underlying processes and is useful for the design of new interventions.

Here, we first show how adversarial training is effective for making deep learning models trained on single-cell RNA-seq data more robust using simulated and real data. Second, we showed that there is a connection between the robustness and interpretability of these deep learning models. Deep learning interpretability methods can identify more significant genes (e.g. markers and key regulators) in cell-type classification tasks when applied to adversarially trained models than the analogous ones without such training. We speculate that adversarial training improves interpretability by constraining gradients to be closer to the data manifold, as seen in other domains (13). We also found that the genes we identify from interpreting our deep learning model are not just the same ones identified by standard differential expression analysis. Presumably, the model is identifying additional important factors in the data to identify these genes and further research is needed to gain a deeper understanding of how this selection process occurs.

Our findings shed light on the important connection between model interpretability and robustness in machine learning. Adversarial training also fortifies a deep learning model, which can be useful for future clinical and health applications, such as diagnostic or prognostic gene expression biomarkers or patient classification, that need to be robust against adversarial attacks (14). We hope our study will encourage researchers to consider adversarial training and the robustness-interpretability relationship in future deep learning research in biology and medicine.

## Method

### Interpretability

We used six interpretability methods to compare our feature importance results:

#### Saliency map

The gradients with regard to inputs *x* are returned by the saliency map as feature importance score *S* (3):

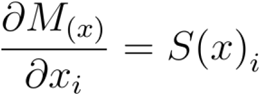

by taking the first-order Taylor expansion of the neural network, M, as in:

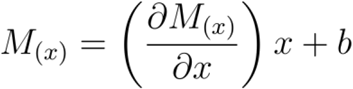

#### Activation maximization

A broad class of methods known as “activation maximization” looks for an input that maximizes the model response, generally using different gradient descent algorithms (19). The idea is to generate an input that best represents the outcome by using:

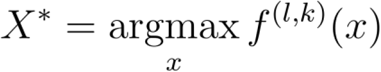

Where x is the input and *X*^*∗*^ is the constructed input which maximizes the activation of a k-th neuron in hidden layer *l* of the neural network *f*.

#### DeepLIFT

By back-propagating, the contribution of every neuron in the network to all features of the input, DeepLIFT decomposes the output prediction of a neural network on a particular input (5) :

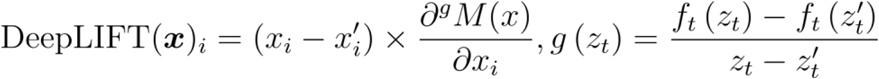

#### DeepSHAP

DeepSHAP calculates the expectations of gradients by randomly selecting baseline data from the distribution and then uses that information to approximate SHAP (SHapley Additive exPlanations) values. Each input sample is first given white noise, and a random baseline is chosen from a predefined distribution. Next, a random point is chosen along the path between the base point and the input with noise, and the gradient of the outputs is computed with respect to the random point. To roughly estimate the expected values, E, of gradients, the technique is done numerous times. The final SHAP score is equal to

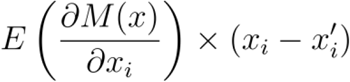

For further description of different interpretability methods implemented in the DeepSHAP package please check (21).

#### LIME

LIME is a surrogate method for explaining the predictions of machine learning models. It generates a new dataset of perturbed samples by making small changes to the original data, then training a simple, interpretable model on this new dataset. It then can be used to explain the prediction of the original model and compute feature importance scores that explain which features of the input had the most influence on the prediction. LIME is formulated as follows:

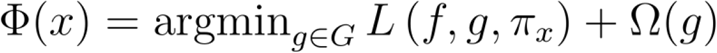

A local model g from class G of interpretable models for an instance x is considered. To avoid having a complex model penalty term, Ω(g) is added. *Πx* denotes the neighborhood of x and L shows the loss between the complex and surrogate model in a defined local neighborhood *Πx*.

### Robustness

In order to generate adversarial data the following methods are used, as found in (41):

#### Fast gradient sign method (FGSM)

Adding practically unnoticeable noise to an input image is a strategy to create an adversarial attack. A common attack method is FGSM (18). FGSM adds noise in the direction of the gradient which reduces the accuracy of the prediction. For an attack level *ϵ*, FGSM sets:

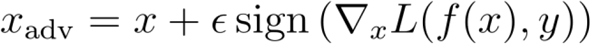

The attack level is chosen to be sufficiently small so as to be undetectable. The optimal ε value depends on the characteristics of the data and specific task.

#### Projected gradient descent (PGD)

PGD is an upgraded version of FGSM that employs several iterations (8). In the equation below, Proj denotes the projection operator, which constrains the input to positions set by a predefined perturbation range. *ε* is the step size with a positive value. PGD works as follows:

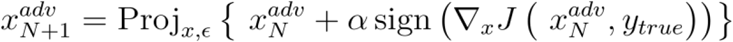

### For visualization, reduced dimension data

Clusters were visualized using the Python package “UMAP”.

## Data preprocessing

The scRNA-seq gene expression data are scaled, centered, and log-normalized. 2000 highly variable genes were chosen. The top 30 principal components were used to map highly variable genes into a lower dimension for clustering. The principal components are further used to generate a k-nearest-neighbour graph with K=30.

## Data availability

We used three scRNA-seq datasets with different characteristics for evaluation. For each dataset, we train a cell-type classification model (the detailed architecture can be found in Supplementary Table 1). Simulated data are generated by SERGIO (17) and can be found at https://github.com/PayamDiba/SERGIO. We downloaded a preprocessed version of the “Development of the murine pancreas” dataset (29) from https://cellrank.readthedocs.io/en/stable/index.html (42). We downloaded a preprocessed version of the “Dentate Gyrus neurogenesis” data (National Center for Biotechnology Information’s Gene Expression Omnibus repository under accession number GSE95753) from https://scvelo.readthedocs.io/en/stable/ (43).

## Code availability

Our Python implementation can be found at:

