## Supplementary material for "Adversarial training improves model interpretability in single-cell RNA-seq analysis": Supp file

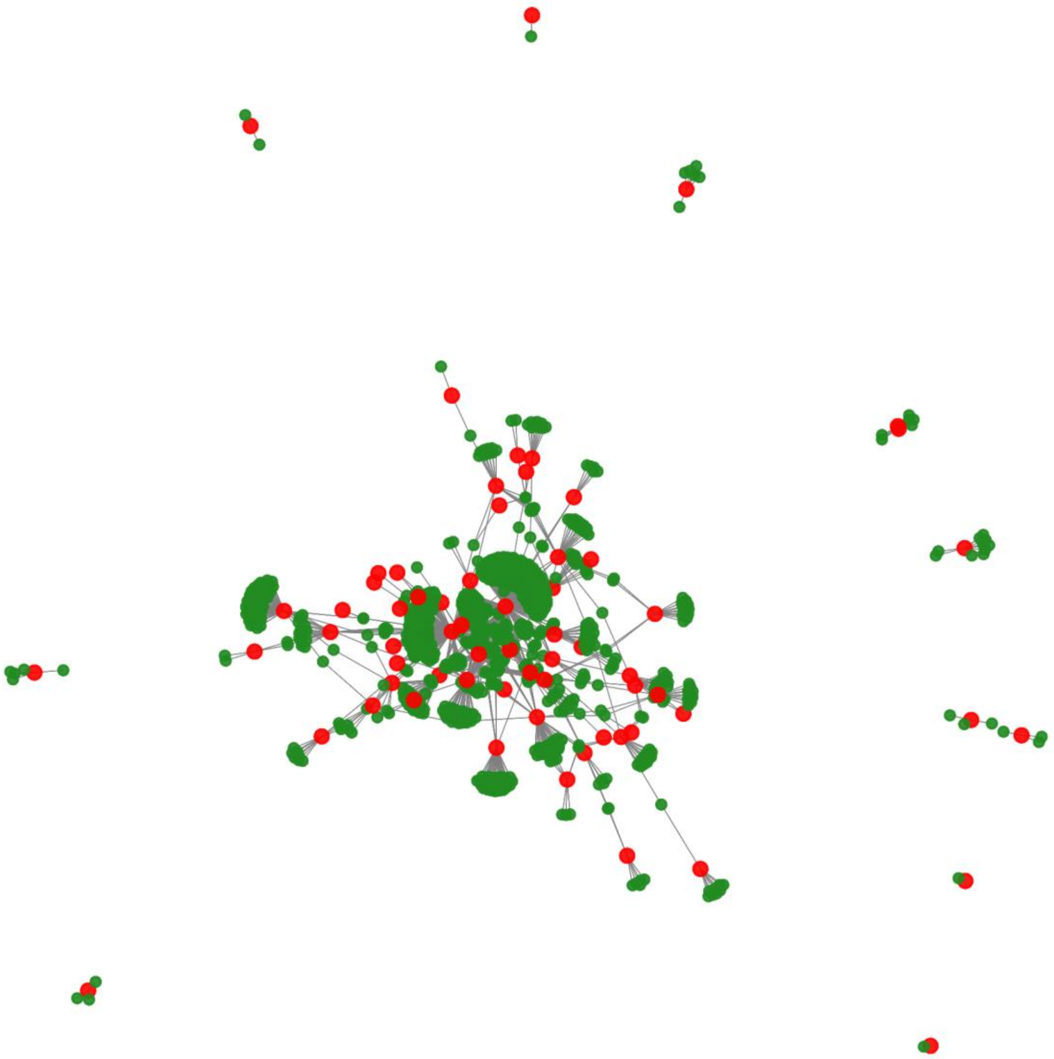

**Supplementary figure 1 - Gene regulatory network of scRNA-seq data for SERGIO.**
The gene regulatory network for SERGIO depicts the interaction between genes within the system. Nodes in red denote key genes (master regulators), while nodes in green represent genes, whose production rates are influenced by their respective regulators.

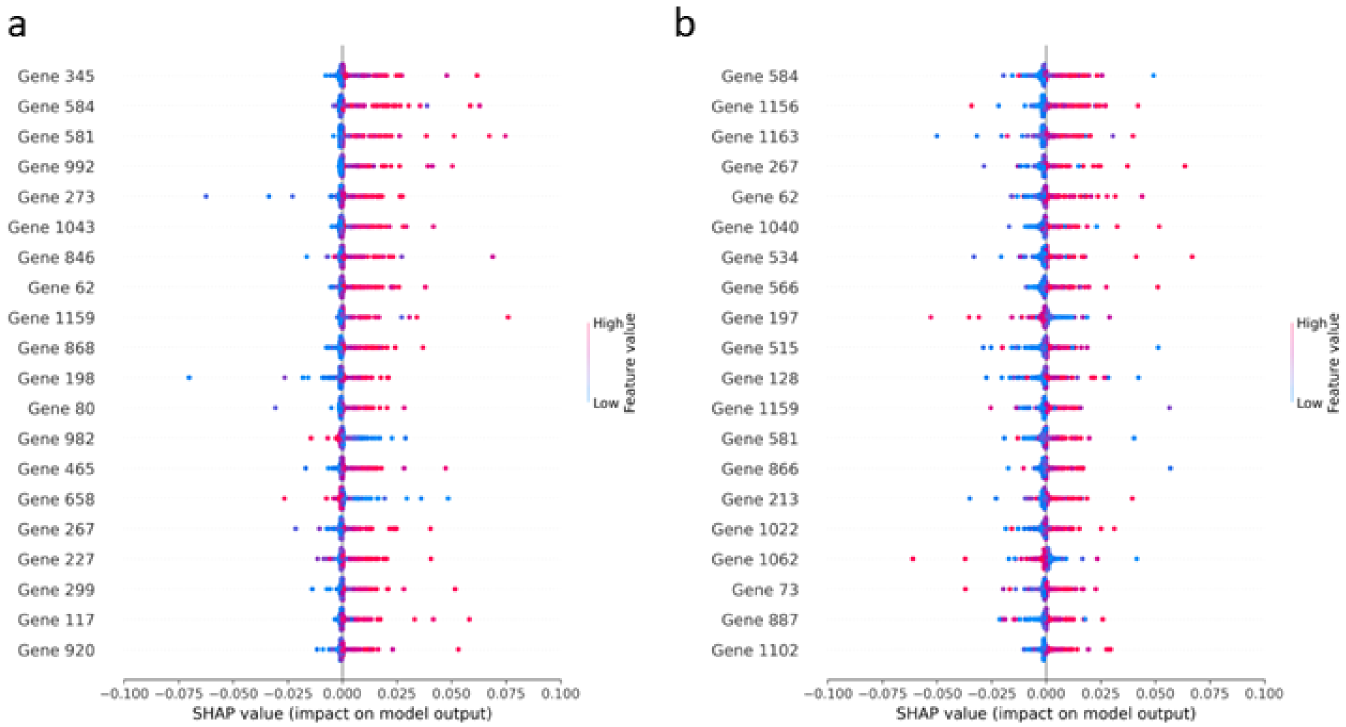


**Supplementary figure 2- Effect of adversarial training on the model's interpretability for the simulated data.**

a, The importance of features (genes) in predicting cell types can be determined by ranking them according to their SHAP values (computed by using Deep explainer) in a standard trained model. b, Ranking genes by using their SHAP values for a model with adversarial training. Each individual data point represents a distinct cell, and the point location on the x-axis indicates the influence of that gene on the model's prediction of the cell-type.

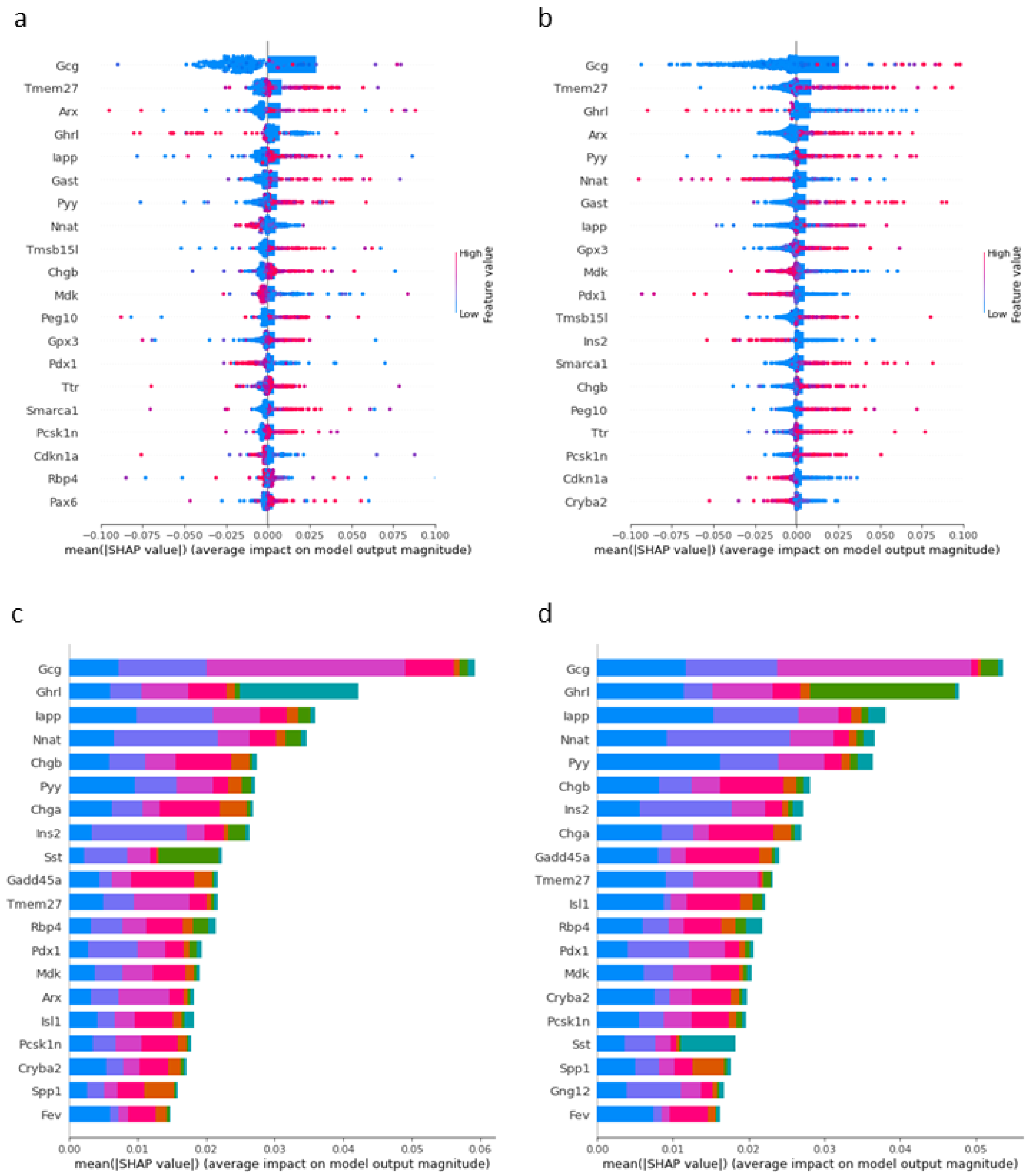


**Supplementary Figure 3: Comparative analysis of different interpretability methods on an adversarially trained model for cell-type classification using mouse Pancreas development data.** (a) and (b) display the top 20 most important genes identified by Deep Explainer and Gradient Explainer, respectively. Each dot represents a gene, and its position on the x-axis corresponds to its SHAP value. (c) and (d) show the mean absolute SHAP values, indicating the rank order of important genes for cell-type classification. These results suggest that different interpretability methods can yield divergent rankings for important genes, highlighting the need for careful consideration of interpretability approaches when analyzing model predictions.

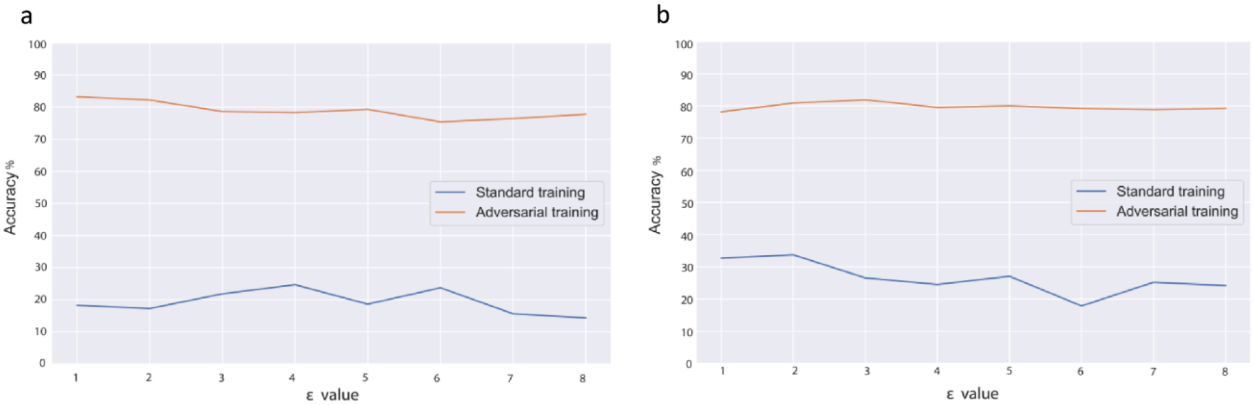


**Supplementary Figure 4: Effect of adversarial attack and training on cell type-classification model accuracy.**

(a) PGD and (b) FGSM methods demonstrate the impact of adversarial perturbation on model accuracy. The orange curves indicate the performance of models with adversarial training, while the blue curves represent models trained using standard methods. The FGSM and PGD attacks have different strengths, and their effect on model accuracy is apparent in the respective subplots. As can be seen, adversarial training significantly improves the robustness of the models against these attacks.

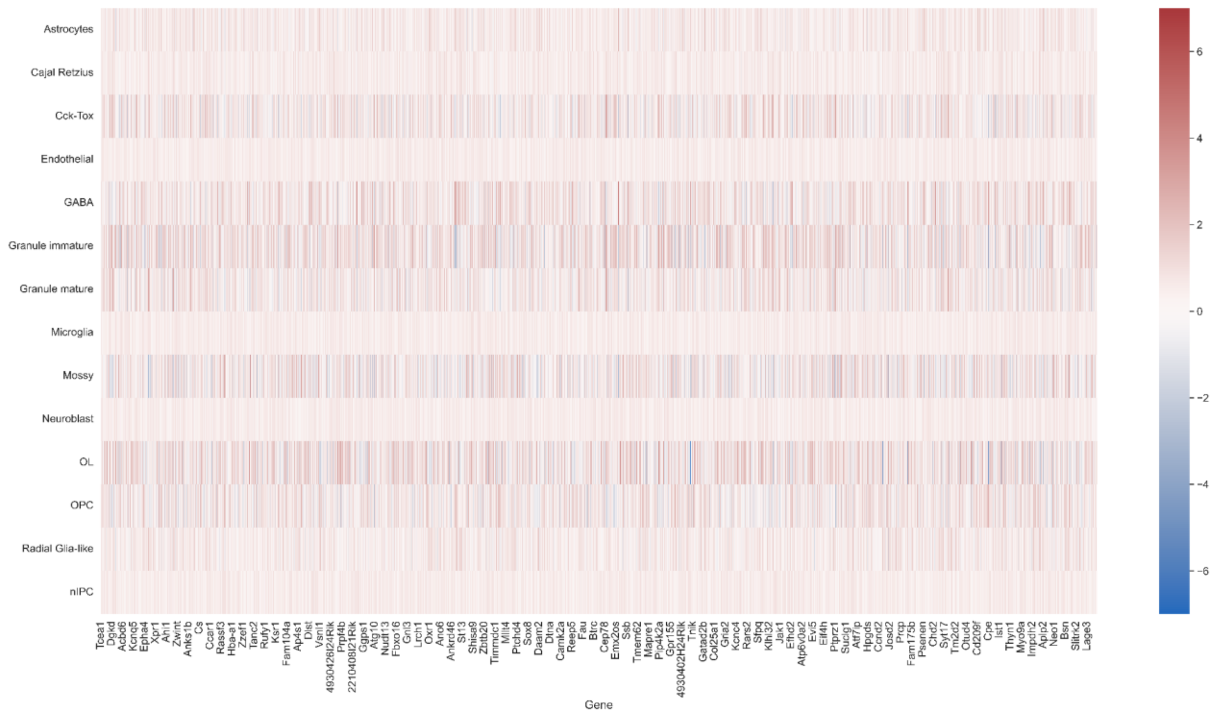


**Supplementary Figure 5 - Activation Maximization scores for each gene-cell type pair for hippocampus development.**

The heatmap of activation maximization scores for gene-cell type pairs. The x-axis corresponds to the genes, while the y-axis represents the cell types. The color of each cell reflects the activation maximization value for the corresponding gene-cell type pair, with red colors indicating higher positive values. This heatmap provides a visual representation of the strength and specificity of gene expression across different cell types, which can help in identifying potential biomarkers or therapeutic targets for specific diseases.

| Name | Type | Size | Activation |
|---|---|---|---|
| Layer 1 | Fully connected | 500 | ReLU and Dropout=0.1 |
| Layer 2 | Fully connected | 250 | ReLU |
| Layer 3 | Fully connected | 100 | ReLU |
| Layer 4 | Fully connected | 80 | ReLU |
| Layer 5 | Fully connected | 60 | ReLU and Dropout=0.2 |
| Layer 6 | Fully connected | 40 | ReLU |
| Layer 7 | Fully connected | 20 | ReLU |
| Layer 8 | Fully connected | Number of cell types | Softmax |
| Optimization: Optimizer: Adam, batch_size:500, epochs: 1000 | | | |

**Supplementary table 1 -** detailed architecture for cell-type classification. The same architecture is used for all the datasets in the paper.
